# Neural oscillatory dynamics reveal altered top-down and integrative mechanisms during face processing in autistic children and unaffected siblings of autistic children

**DOI:** 10.1101/2025.11.03.684159

**Authors:** Theo Vanneau, Chloe Brittenham, Megan Darrell, John J. Foxe, Sophie Molholm

## Abstract

Face processing is fundamental to social communication and has been a major focus of autism research. While event-related potential (ERPs) studies of face processing have produced mixed results, little work has examined neuro-oscillatory dynamics, which may better capture the integrity of underlying networks. To address this gap, EEG was recorded from children aged 8-13 across three groups: autistic (*n* = 50), non-autistic (*n* = 38) and siblings of autistic children (*n* = 26), during a visual oddball task. In a blocked design, participants viewed faces and objects, presented upright and inverted (non-targets), to assess the face inversion effect (the FIE; a larger or earlier N170 to inverted than upright faces), and responded to infrequent shadow versions (targets). Analyses using permutation statistics and linear mixed models focused on non-target stimuli, quantifying face-related ERPs (P1, N170) and oscillatory activity associated with sensory and attentional processing (theta, alpha, gamma). Across groups, faces elicited earlier P1 and larger N170 amplitudes than objects, and showed a FIE. Furthermore, the rightward lateralization of the FIE was reduced for autistic participants. Analyses in the frequency domain revealed greater induced theta for inverted versus upright stimuli and for faces versus objects, revealing face specific effects, and stronger theta for inverted faces for the autistic and sibling groups, suggesting greater cognitive effort in processing these social stimuli. Gamma-band inter-trial phase coherence exhibited face selectivity only in the non-autistic group, pointing to differences in early network synchronization in autistic children relative to their non-autistic peers, whereas alpha event-related desynchronization did not vary by group or category. Altogether, these findings support altered neural synchronization/efficiency for autistic participants and siblings of autistic children, that is specific to face stimuli and seen despite largely typical sensory driven encoding. These data suggest that neural oscillatory assays are more sensitive to face processing differences in autism than broadband ERPs and that these oscillatory assays may be endophenotypic.

## Introduction

### Tuning to Faces

Faces are among the most socially and behaviorally salient visual stimuli, and the ability to rapidly and accurately process and interpret them is fundamental to human social functioning. Facial cues convey essential information about identity, emotional state, gaze direction, and communicative intent; elements that must often be decoded within milliseconds to guide appropriate behavioral responses. The human brain begins to differentiate faces from other visual stimuli remarkably quickly, with face-selective neural activity detectable as early as ∼100 milliseconds post-stimulus onset (Colombatto & McCarthy, 2017; Rousselet et al., 2007). Consistent with this specialization, behavioral studies demonstrate that faces are detected more rapidly than control stimuli such as houses (Purcell & Stewart, 1986) and this detection advantage appears to widen across development (Tottenham et al., 2006). These rapid responses reflect a highly specialized and temporally efficient face processing system, shaped by both bottom-up sensory inputs and top-down modulatory influences such as attention, expectation, and experience (Hadders-Algra, 2022). This system involves a distributed network of cortical and subcortical regions, including the fusiform face area, occipital face area , superior temporal sulcus, and amygdala, each contributing to distinct aspects of face perception (Haxby et al., 2000; Johnson, 2005; Johnson et al., 2015; Kanwisher et al., 1997).

Event-related potentials (ERPs) have long been a primary tool for probing the temporal dynamics of face perception, offering high temporal resolution and a well-established set of components that index stages of face-selective processing. The P1 component, peaking around 100 ms at occipital electrodes, reflects early visual processing and is often modulated by the presence of faces (Olivares et al., 2015). The N170, a negative deflection observed around 170 ms maximally over lateral occipito-temporal regions, is a robust electrophysiological signature of face perception (Keyes et al., 2010). The sensitivity of N170 to faces over non-face objects has been extensively documented (Allison et al., 1994; Itier et al., 2011; Rossion & Jacques, 2011; Rossion et al., 2012; Yovel, 2016). Furthermore, N170 has been consistently shown to be larger and delayed for inverted faces compared to upright faces, commonly referred to as the face inversion effect (FIE; Bentin et al., 1996; Botzel et al., 1995; Itier et al., 2006; Jeffreys, 1989; Rossion et al., 1999).

### Face processing in autism

Autism spectrum disorder (autism) is a neurodevelopmental condition characterized by differences in social communication and interaction, alongside the presence of restricted interests and repetitive behaviors (American Psychiatric Association, 2013). Among the most consistently reported findings in autism are atypical responses to faces and facial expressions, which have been suggested to contribute to broader difficulties in social cognition (Dawson et al., 2004; Kliemann et al., 2010; Senju, 2013). These atypicalities often include reduced spontaneous attention to faces, diminished eye contact, and difficulties interpreting emotional expressions (Chawarska et al., 2013; Constantino et al., 2017; Harms et al., 2010; Senju & Johnson, 2009). Eye-tracking studies have shown that autistic individuals tend to exhibit altered scanning patterns, such as fewer fixations to the eye region and reduced overall face engagement (Pelphrey et al., 2002). Behavioral studies report difficulties in face identity recognition and emotion discrimination in autistic individuals, with meta-analyses suggesting moderate to large effect sizes for challenges in face identity processing in autism (Griffin et al., 2023), although performance can be similar to their non-autistics peers in certain contexts (see Tang et al., 2015; Weigelt et al., 2012 for reviews). Notably, behavioral effects are often more robust than corresponding neural differences, raising important questions about the timing and nature of face processing differences in autism (Griffin et al., 2023). Electrophysiological studies using ERPs illustrate this point, with components such as the N170 producing inconsistent results in autism. A comprehensive review found that among 23 ERP studies investigating the N170 component in autism, the majority reported no significant group differences in amplitude (18/23 studies) or latency (17/23 studies) in response to faces (Feuerriegel et al., 2015). More recent work similarly suggests broadly typical early perceptual mechanisms, (Aydin et al., 2023; Shephard et al., 2020), although some studies report delayed N170 latencies or attenuated amplitudes in autistic individuals, possibly reflecting subtle differences in early visual encoding (Deng et al., 2025a).

Importantly, some effects that are robust at the behavioral level, such as the FIE (behaviorally characterized by lower accuracy/sensitivity and longer reaction times), are less consistently observed in ERPs. A recent meta-analysis combining behavioral and ERP data found that FIE is reduced in autistic individuals compared to neurotypical peers. This reduced sensitivity is hypothesized to reflect a weaker specialization of the face processing system, with greater reliance on featural analysis, particularly under conditions of increased cognitive demand (Griffin et al., 2023). This pattern suggests that face processing differences in autism may emerge more clearly at later processing stages involving memory, integration, or attentional control. Taken together, these findings point toward a model in which early, automatic neural responses to faces may be largely typical in autism, but differences emerge at stages of processing that require more effortful or integrative operations.

In non-autistic individuals, face processing usually shows right-hemispheric dominance (e.g., greater N170 amplitude over right occipitotemporal sites; Bentin et al., 1996). In autism, studies report reduced lateralization, more bilateral patterns, or even left-dominant responses (Deng et al., 2025a, 2025b). Reduced lateralization may reflect differences in neural organization and/or the engagement of compensatory mechanisms. Investigating asymmetry may provide insight into developmental pathways and processing strategies, and atypicalities in hemispheric lateralization for object category processing have been previously described by our group using high-density EEG (Fiebelkorn et al., 2013).

### Oscillatory Dynamics in Face Perception

Time-domain ERP components provide only a partial picture of the neural processes involved in face processing and perception. By averaging across trials and retaining only phase-locked activity, ERPs potentially obscure important non-phase-locked responses and collapse the diversity of underlying neural mechanisms into a handful of experimenter-defined peaks. Although statistical techniques such as permutation testing can mitigate some subjectivity in component selection, ERPs ultimately reflect the summed output of many processes rather than their distinct contributions.

Oscillatory dynamics provide an expanded window into how neural populations coordinate activity during face perception (Guntekin & Basar, 2014). Spectral analyses decompose the EEG signal into its constituent frequencies revealing oscillatory activity that is not readily observable in time-domain data alone. Event-related changes in theta (4-7Hz), alpha (7-13Hz), and gamma-band (25-40Hz+) activity have been directly linked to specific perceptual and cognitive operations, offering a more mechanistically grounded approach. In neurotypical individuals, these rhythms index processes supporting face encoding and attention. Among these rhythms, increased gamma activity has been tied to the integration of facial features into coherent representations (Balconi & Lucchiari, 2008; Sato et al., 2014; Zion-Golumbic & Bentin, 2007) with gamma intertrial phase coherence (ITPC) reflecting the temporal alignment of neural responses. Although not commonly assessed, there is some support for reduced gamma activity in response to faces in autism (Dawson et al., 2005; Sun et al., 2012).

Alpha activity is widely understood to implement an inhibitory gating mechanism; its event-related desynchronization (alpha ERD) marks a release of that inhibition and thus indexes increased cortical excitability within task-relevant networks. In this view, the magnitude of alpha ERD reflects a joint function of stimulus features and the degree of attentional engagement directed toward the stimulus (Hanslmayr et al., 2011; Klimesch et al., 2007; Pfurtscheller & Klimesch, 1992). More broadly, alpha rhythms coordinate attentional selection, sensory inhibition, and excitability control (Foxe et al., 2014; Foxe et al., 1998; Foxe & Snyder, 2011; Klimesch, 1999; Snyder & Foxe, 2010). To our knowledge, alpha ERD to faces has not yet been systematically examined in autism, although our group has shown dysregulation of alpha-band suppression mechanisms in autism in the context of intersensory selective attention (Murphy et al., 2014), suggesting that this is a candidate mechanism for further investigation in this population.

Theta-band activity is associated with cognitive control, working memory, and large-scale network coordination. Frontal event-related theta indexes control and effort (Cavanagh & Frank, 2014) and varies with face familiarity (Basar et al., 2007; Zion-Golumbic et al., 2010), species specificity (conspecific faces vs. objects in non-human primates; (Zhang et al., 2024)), and emotional content (Balconi & Lucchiari, 2006; Guntekin & Basar, 2014). During face recognition, intracranial recordings have shown robust theta–gamma coupling in the inferior occipital gyrus, suggesting that theta rhythms help orchestrate gamma-band activation (Sato et al., 2014). Developmentally, theta-band face specificity is reduced in young autistic children (Dawson et al., 2012) and theta power increases with socially focused intervention in children at elevated likelihood for autism (Dawson et al., 2012; Jones et al., 2017). In sum, different neural oscillatory frequency bands are engaged for coordinated functions related to face processing, but to date these have not been extensively characterized in autism.

### Broad Autism Phenotype

A key question concerns whether the neural differences observed in autism reflect disorder-specific alterations or instead index broader familial traits. Siblings represent a genetically informative comparison group: while they do not meet diagnostic criteria for autism, they often exhibit subclinical features or neural traits associated with the broader autism phenotype (BAP; Ingersoll & Wainer, 2014; Losh et al., 2009; Losh & Piven, 2007; Pisula & Ziegart-Sadowska, 2015). Prior studies have shown that SIBs may display intermediate behavioral and neurophysiological profiles between non-autistic and autistic individuals, particularly in domains such as sensory responsiveness and attentional control (Matthews et al., 2013; Narvekar et al., 2022; Pani et al., 2013). This is especially relevant for face processing and social communication in which alterations are increasingly recognized as early-emerging and heritable components of the autistic phenotype (Constantino et al., 2017; de Klerk et al., 2014; Losh & Piven, 2007). Therefore, the current study also included a group of unaffected siblings (SIBs) of children with autism allowing us to assess whether any neural markers of face processing differences may reflect familial traits potentially linked to genetic liability.

### The Current Study

We employed high-density EEG to investigate neural responses to social and non-social stimuli in autistic children, unaffected siblings of autistic children, and non-autistic controls. Participants completed a visual oddball task while EEG data were recorded to assess event-related potential (ERPs) and neural oscillatory indices of face processing, sensory integration, and attentional control. We hypothesized that, in line with prior work, ERP markers of face processing (P1, N170) would be largely similar among groups. In contrast, we expected differences to more clearly emerge in oscillatory dynamics, particularly reduced gamma selectivity for faces in autism. Finally, we predicted that siblings would exhibit intermediate neural profiles between non-autistic and autistic groups, consistent with these representing genetically transmitted differences in neural functions.

## Methods

### Participants

The study initially included 42 non-autistic (NA), 61 autistic (AU), and 27 sibling (SIB) participants, all between 8 and 13 years of age. After exclusions based on behavioral, EEG and eye-tracking data quality (see corresponding sections for data exclusion criteria), final analyses included 38 NA, 50 AU, and 26 SIB participants (see Table 1 for participant characteristics). To be included in the AU group, participants were required to meet diagnostic criteria for autism spectrum disorder on the basis of the following measures: 1) Autism Diagnostic Observation Schedule, Second Edition (ADOS-2; Lord et al., 1994), diagnostic criteria for autistic disorder from the *Diagnostic and Statistical Manual of Mental Disorders* (DSM-5; American Psychological Association, 2013), and clinical impression of an experienced licensed clinician. Due to COVID-19 precautions, a subset of AU participants (*n* = 9) could not complete the ADOS-2 evaluation as masking requirements impacted administration and instead underwent the Childhood Autism Rating Scale 2 (CARS-2) and Autism Diagnostic Interview-Revised (ADI-R) for diagnostic assessment. NA participants met the following inclusion criteria: no history of neurological, developmental, or psychiatric disorders, no first-degree relatives diagnosed with autism, and enrollment in an appropriate age grade in school. The SIB group participants met the same criteria as the NA group, except for having a first-degree biological sibling diagnosed with autism. Exclusion criteria across groups included: (1) a known genetic syndrome associated with an IDD (including syndromic forms of autism), (2) a history of or current use of medication for seizures in the past 2 years, (3) significant physical limitations (e.g., vision or hearing impairments, as screened over the phone and on the day of testing), (4) premature birth (<35 weeks) or having experienced significant prenatal/perinatal complications, or (5) a Full Scale IQ (FS-IQ) of less than 80. All procedures were approved by the Institutional Review Board of the Albert Einstein College of Medicine and adhered to tenets for human subjects’ research laid out in the Declaration of Helsinki. All participants assented to the procedures and parents/guardians provided informed consent. Participants received nominal recompense for their participation (at $15 per hour).

**Table 1.**
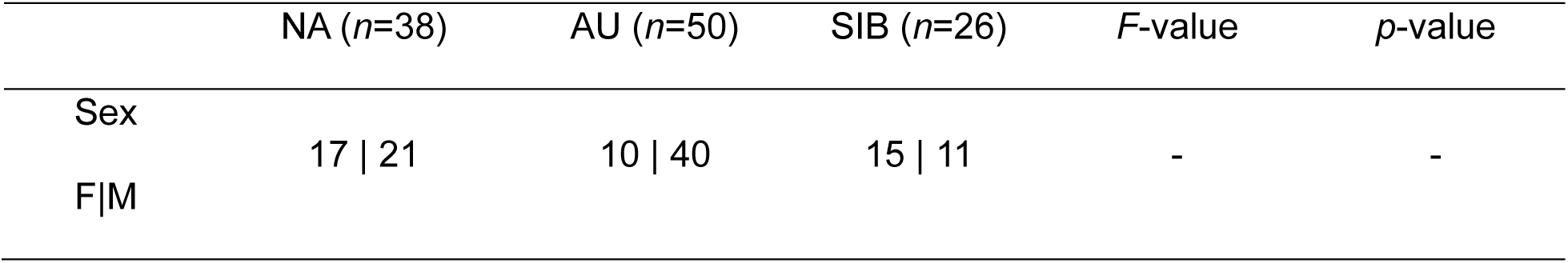
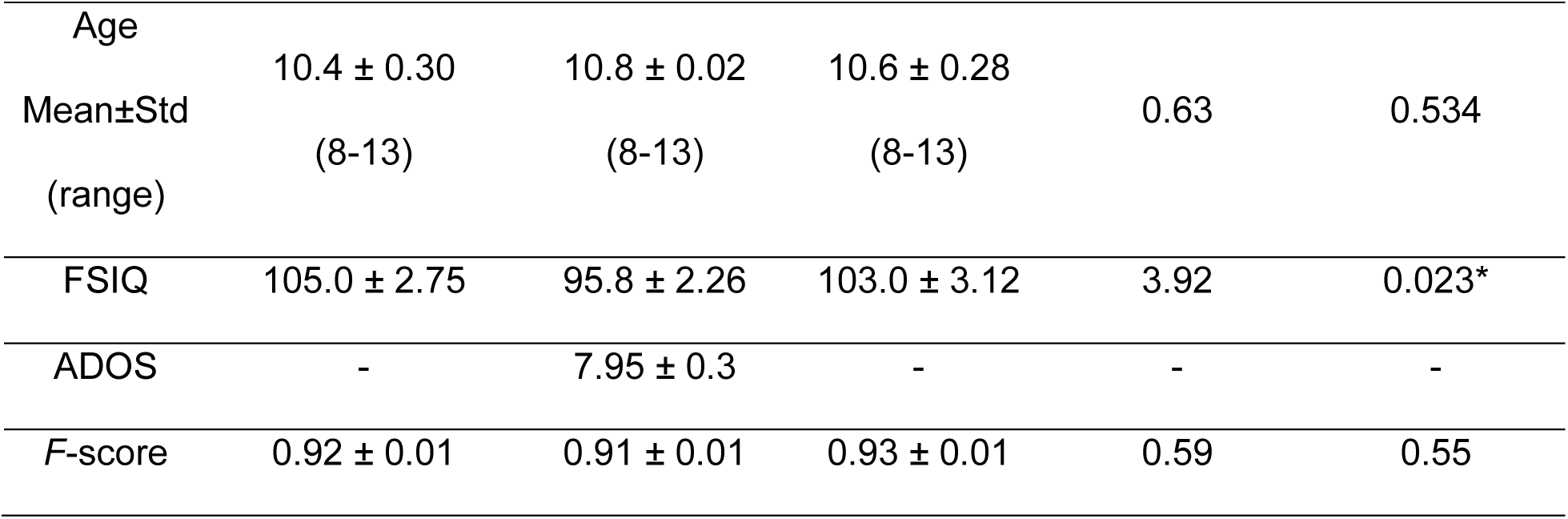
Demographic characteristics.

### Experimental procedures

Participants were seated in a chair in an electrically shielded room (International Acoustics Company, Bronx, New York), 70 cm away from the visual display (Dell UltraSharp 1704FPT). The stimuli, controlled by Presentation software (Neurobehavioral Systems), were faces (’Social’) or objects (‘Non-Social’), each shown as upright and inverted images, along with infrequently presented shadow versions. The faces come from the NimStim database (Tottenham et al., 2009) and the objects from the BOSS database (Brodeur et al., 2010). Participants were instructed to press a button as quickly as possible upon detecting a shadow stimulus (presented at 20% probability, see Figure 1A for illustration). A jittered interstimulus interval (900–1100ms) reduced onset predictability. The task comprised 720 trials across 12 blocks (60 trials/block, ∼3 minutes and 40 seconds each). Blocks were organized by stimulus category; each block contained only social stimuli (i.e., upright and inverted faces with their shadow versions) or non-social stimuli. There were six blocks of social and six blocks of non-social stimulus in total. Face and object images (upright/inverted) were randomly chosen from a pool of 28 stimuli. All faces depicted a positive emotion, (i.e., smiling faces). Shadow faces and objects were chosen across a reduced pool of 5 stimuli. Responses were recorded using a response pad (Logitech Wingman Precision Gamepad), and stimulus and response triggers were sent from the PC acquisition computer via Presentation software. See Figure 1A for an illustration of the experimental paradigm.

**Figure 1.**
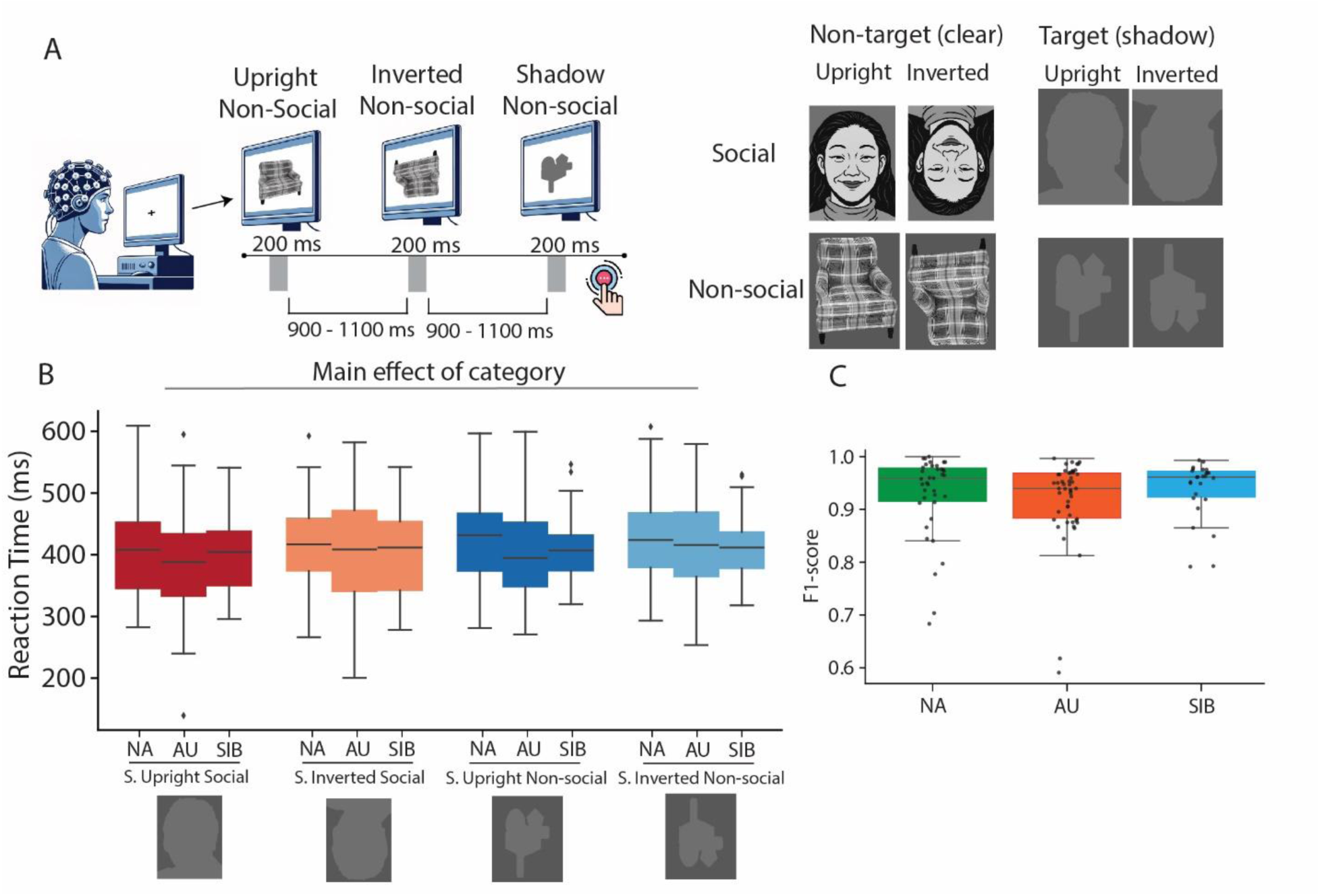
Paradigm design and behavioral performance. **(A)** Schematic of the paradigm with the 8 different possible stimulus types (note that social and non-social stimuli are presented in separate blocks) where the participants are tasked to respond to shadow stimuli. Social stimuli were real face images (NimStim database); for display purposes here, we show stylized comic representations in adherence with the database stipulations. (**B**) Reaction time of each group to each type of shadow stimulus (reported main effect by using LMMs) and (**C**) corresponding F1-score.

### Behavioral data analysis

To ensure data quality, an initial behavioral filtering step was applied. The following performance metrics were calculated for each participant: 1) Miss rate: proportion of missed responses (*false negatives / total stimuli*), 2) False alarm: proportion of responses to non-target stimuli (*false positives / total stimuli*), 3) Precision: accuracy of correct responses (*true positives / (true positives + false positives*)), 4) Recall: proportion of correct detections (*true positives / (true positives + false negatives*)), and 5) F-score: combined measure of precision and recall (*2 × (precision × recall) / (precision + recall*)). Based on these criteria, six participants from the AU group were excluded due to non-compliance with the task, defined as having either a false alarm rate exceeding 50% (among them 2 AU participants were responding to every stimulus) or a Miss rate exceeding 50% (which included 1 AU participant who was never responding).

### Eye-tracking recordings

Gaze position and pupil data (not reported here) were recorded using EyeLink 1000 (SR Research Ltd., Mississauga, Ontario, Canada) at a sampling rate of 500 Hz. Two NA, five AU and one participant in the SIB group were excluded due to the absence of eye-tracking recordings caused by hardware or measurement issues (e.g., interference from glasses, eyelid occlusion). To further refine trial selection, gaze position data were used to filter all trials, with trials rejected when the participant was not fixating within a defined region of interest centered on the fixation cross (±7° from fixation in the x-axis and ±5° on the y-axis). This filtering was applied over a 40 ms window around stimulus onset (−20 ms to +20 ms). The proportion of trials rejected due to eye-tracking filtering was as follows: NA group: 6.27% ± 1.0, AU group: 15.94% ± 2.45; SIB group: 6.84% ± 1.68 (Mean ± SEM).

### EEG recordings & preprocessing

EEG data were recorded at a sampling rate of 512Hz using 64 channels from a BioSemi Active II system (using the CMS/DRL referencing system) with an anti-aliasing filter (−3 dB at 3.6 kHz). Analyses were conducted in Python (3.11) using MNE (Gramfort et al., 2013) and custom scripts available at https://github.com/tvanneau/SFARI-FAST. Bad channel detection was performed using the function NoisyChannels (with RANSAC) from the pyprep toolbox (Bigdely-Shamlo et al., 2015). If more than 15% of the channels were detected as bad, the participant was rejected (1 NA, 1 SIB). Bad channels were interpolated using spline interpolation (Perrin et al., 1989). EEG was filtered using a FIR band-pass filter (0.01-40Hz), and Independent Component Analysis (ICA) on 1Hz high-pass EEG (just for the ICA analysis) was used to identify and manually reject eye-related components (blinks/saccades). Epochs were created from -500 to +1000ms with a baseline-correction from -200ms to -50ms relative to stimulus onset. All analyses were performed on the responses to non-target stimuli. For all analyses, EEG epochs were referenced to a common average reference.

### ERP analysis

Amplitudes and latencies of the face sensitive P1 and N1 were extracted by identifying the local maximum value for the P1 or minimum value for the N170 within the time ranges of 50–180 ms and 170–200 ms respectively, from channels over central occipital scalp (‘O1’ and ‘O2’) where the P1 response was maximal (Aydin et al., 2023; Colombatto & McCarthy, 2017; Foxe & Simpson, 2002; Shephard et al., 2020), and from channels over lateral-occipital scalp (‘P7’ for the left hemisphere and ‘P8’ for the right hemisphere) where the N170 was maximal (Aydin et al., 2023; Doniger et al., 2000; Doniger et al., 2001), respectively. The broader P1 window was acceptable given its sharp, highly consistent peak across individuals, reflected in low latency variability, whereas a narrower window for the N170 was necessary to differentiate the N170 response from a subsequent negative component at about ∼250 ms. In addition to the above electrode and temporally constrained analyses, permutation statistics on difference waves were applied to further investigate the FIE (using the difference between the ERP to upright versus inverted faces).

### Spectral analysis

Spectral analyses were performed using complex Morlet wavelet convolution (tfr_morlet function in MNE) between 2 and 40Hz with a Full Width at Half Maximum (FWHM) in the spectral domain of 3.25Hz and 187ms in the temporal domain (Cohen, 2014) that correspond to: number of cycles = frequency / 2 (with frequency as a vector of frequency from 2– 40 in steps of 0.19Hz). The non-phase-locked power was calculated after subtracting the stationary ERP from each trial for each individual (Kalcher & Pfurtscheller, 1995). For the total power and the non-phase-locked power, an average baseline normalization was applied after averaging all the trials using a -200ms to 0ms baseline period and power was expressed as percentage of change relative to that baseline. Analyses focused on the theta (4–7Hz), alpha (7–13Hz) and low gamma (25–40Hz) frequency bands.

## Statistical analysis

Analyses were conducted using Jamovi (The jamovi project, version 2.3.28, https://www.jamovi.org for analysis of Variance (ANOVA) and linear mixed models (LMMs)) and in Python (MNE-Python) for permutation-based statistics (Maris & Oostenveld, 2007)). Group differences in Age, FSIQ and F-score were assessed using ANOVA (α = 0.05) with group as a between-subjects factor after confirming normality using a Shapiro-wilk test (Shapiro & Wilk, 1965). Between-groups comparisons of RTs, latency and amplitudes of the P1 and N170, induced theta power and gamma ITPC were performed using LMMs to account for individual variability, with fixed factors of Group (NA, AU, SIB), Orientation (upright, inverted), Category (social, non-social), and Hemisphere (right, left), with age as a covariate; all interactions were tested. Post-hoc tests were Bonferroni corrected. Within each group, the FIE was tested with nonparametric permutation methods (Maris & Oostenveld, 2007) on the ERP difference wave (Inverted-social − Upright-social) using two complementary approaches: (i) a region-of-interest test over preselected temporal channels (P7 and P8) to confirm the FIE, implemented as one-sample t-tests against zero with t_max_ (max-T) correction; and (ii) an exploratory spatio-temporal cluster-based permutation test without preselecting channels or time windows, again using one-sample tests against zero. For oscillatory power and gamma ITPC, the channels were selected based on topographical activity and we applied temporal cluster-based one-sample tests. All permutation procedures used 1,000 permutations (α = 0.05), comparing observed t-values to null distributions of maximal t-statistics obtained from label shuffling. Correlations were computed with Pearson’s *r* when data were normally distributed and Spearman’s *ρ* otherwise, with normality assessed via the Shapiro–Wilk test.

## Results

### Faster Reaction Times to Faces Across All Groups

Across all groups, participants responded faster to social than non-social target stimuli and faster to upright than inverted target stimuli (Category: *F*(1,330) = 28.96, *η²p* = 0.08, *p* < .001, Δ = 15.9 ms; Orientation: *F*(1,330) = 6.21, *η²p* = 0.018, *p* = .01, Δ = 7.38 ms; Figure 1B). Older participants responded more quickly (*F*(1,109) = 15.53, η²p = 0.12, *p* < .001; estimate: -15.9 ms/year). There was no main effect of group or group by category interactions (*p*s > .53). F-scores did not differ significantly across groups (*p* > .50; Figure 1C).

### Early visual ERP measures: P1 and N170

The visual evoked potentials (VEPs) in response to all stimulus types and for all groups consisted of a prominent positive going response over posterior occipital scalp, the P1, followed by a negative going (but still positive) bilaterally focused response over parieto-temporal scalp regions, the N170 (Figure 2A; Figure 3A). This pattern is consistent with VEPs typically observed at this stage of development (Altschuler et al., 2014; Knight et al., 2024; Kuefner et al., 2010).

**Figure 2.**
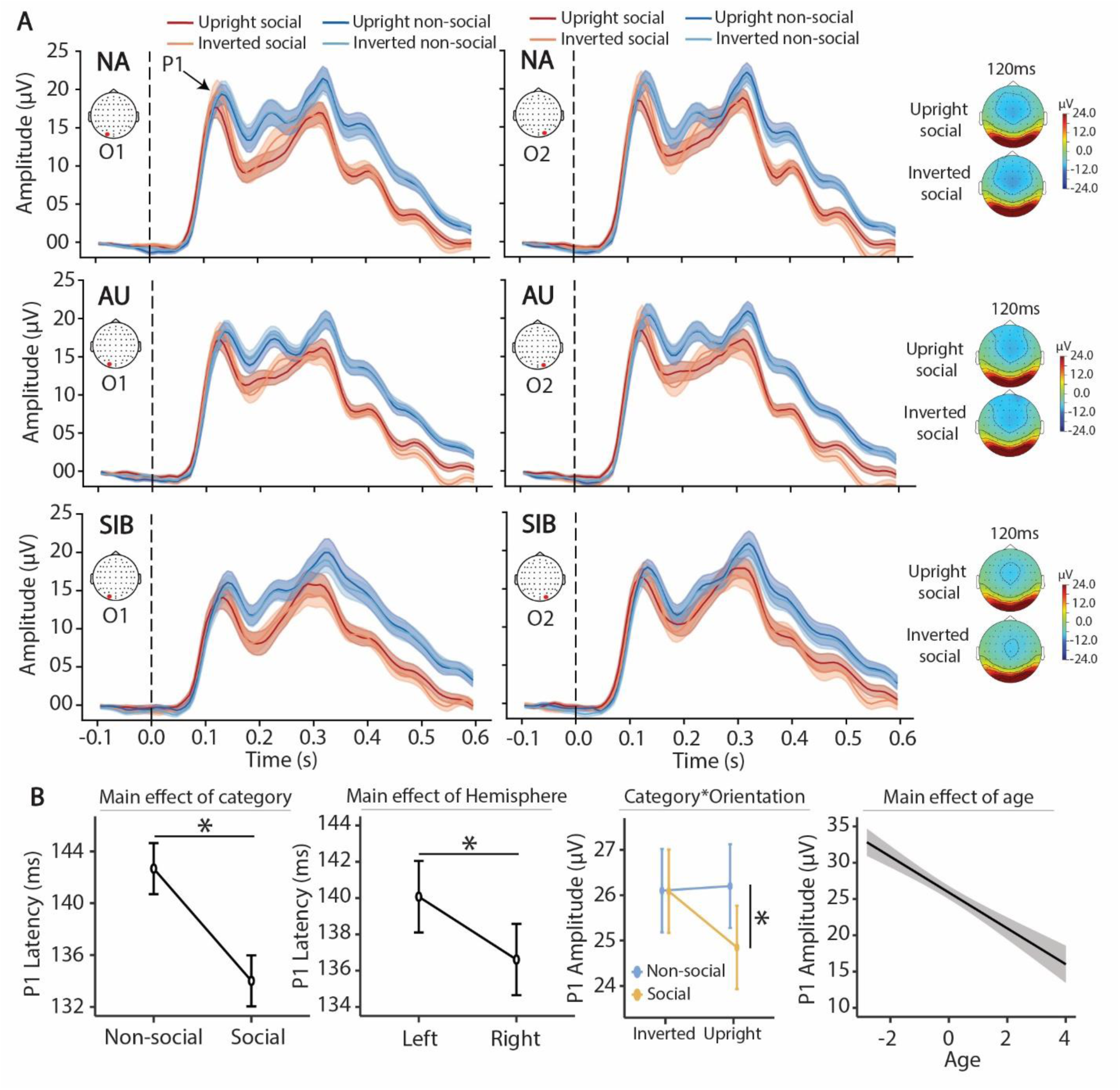
P1 responses to each stimulus type by group. **(A)** ERP measured over O1 (left-side) and O2 (right-side) for non-autistic (NA; upper row), autistic (AU, middle row) and siblings of autistic children (SIB; bottom row) in response to Upright social (red), Inverted social (orange), Upright non-social (blue) and Inverted non-social (light blue) with associated topographical map showing the activity at the timing of the P1 (120 ms) in response to Upright social and Inverted social. **(B)** Significant main effects revealed by the LMMs: Latency of the P1 (latency of the peak within 80–160 ms) for each category; P1 latency for the left and right hemisphere; P1 Amplitude by category and orientation; Effect of age on P1 amplitude. (*) indicates significant post-hoc (α = 0.05 and Bonferroni condition).

**Figure 3.**
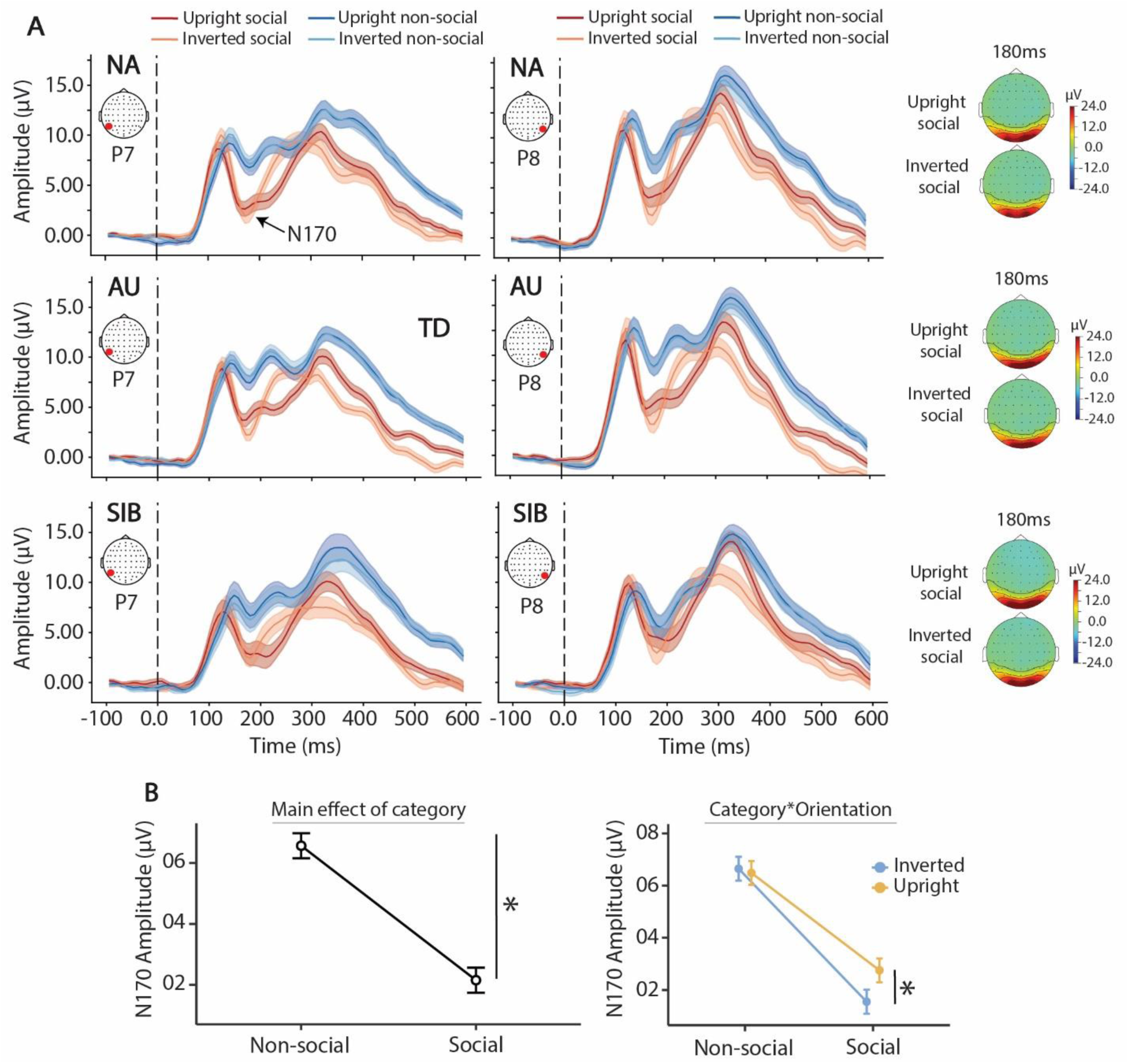
N170 responses to each stimulus type by group. **(A)** ERP measured over P7 (left-side) and P8 (right-side) for non-autistic (NA; upper row), autistic (AU, middle row) and siblings of autistic children (SIB; bottom row) in response to Upright social (red), Inverted social (orange), Upright non-social (blue) and Inverted non-social (light blue) with associated N170 topographical map in response to Upright social and Inverted social. **(B)** Significant main effects revealed by the LMMs: N170 amplitude for each category and orientation by category interaction. (*) indicates significant post-hoc (α = 0.05 and Bonferroni condition).

### P1 Latency

LMMs revealed that P1 peak latency was earlier to social compared to non-social stimuli (Figure 2B; Category: *F*(1,770) = 46.39, *η²p* = 0.06, *p* < .001; Δ = 8.66 ms; Δ = 0.68 µV), and earlier over the right hemisphere (Hemisphere: *F*(1,770) = 7.43, *η²p* = 0.01, *p* = .007; Δ = 3.47 ms). A significant Category by Orientation interaction (*F*(1,770) = 8.07, *η²p* = 0.01, *p* = .005) reflected that P1 peak-latency was earlier to upright compared to inverted non-social stimuli (Δ = 4.90 ms, *t_770_* = 2.73, *p* = .039) whereas it did not differ between upright and inverted social stimuli (*p* = 1.0).

### P1 Amplitude

P1 amplitude was greater for non-social than social stimuli (*F*(1,770) = 7.01, *η²p* = 0.01, *p* = .008) and for inverted compared to upright stimuli (*F*(1,770) = 4.86, *η²p* < 0.01, *p* = .028; Δ = 0.56 µV). Further, P1 amplitudes were larger over the right hemisphere (*F*(1,770) = 18.33, *η²p* = .02, *p* < .001; Δ = 1.11 µV), an effect driven by the SIB group (Group*Hemisphere: *F*(2,770) = 5.24, η²*p* = 0.01, *p* = .005), with a significant *post-hoc* test between left and right hemisphere only for the SIB group (Δ = 2.29 µV, *t_770_* = 4.41, *p* < .001). An Orientation by Category interaction (*F*(1,770) = 6.71, η²*p* = 0.01, *p* = .01) reflected higher amplitude for inverted than upright social stimuli (Δ = 1.23 µV, *t_770_* = 3.39, *p* = .004). LMM also revealed a main effect of age (*F*(1,109) = 19.61, η²*p* = 0.15, *p* < .001) with P1 amplitude decreasing for older participants (Estimate: -2.44 µV/year). See Figure 2B.

### N170 Latency

LMMs revealed a significant effect of age (*F*(1,109) = 19.80, η²*p* = 0.154, *p* < .001) with earlier N170 for older participants (*ß* = 1.59 ms, *t* = 4.45), and a significant effect of Orientation (*F*(1,770) = 10.02, η²*p* = .013, *p* = .002) with earlier N170 for inverted stimuli (Δ = 1.60 ms, *t*_770_ = 3.17, *p* = .002). Other main effect comparisons (Group, Category, Hemisphere) were not significant (*p*s > .11).

### N170 Amplitude

LMMs revealed a significant main effect of Category (Figure 2E; *F*(1,770) = 240.36, η²*p* = 0.24, *p* < .001) with larger N170 amplitude for Social compared to Non-social stimuli (Δ = 4.41 µV, *t* = 15.5, *p* < .001). The Orientation by Category interaction was also significant (*F*(1,770) = 5.74, η²*p* = 0.01, *p* = .01), with *post-hoc* testing showing a larger N170 for inverted compared to upright, for social stimuli only (Δ = 1.20 µV, *t* = 2.98, *p* = .018). See Figure 3B. This analysis did not reveal interactions with Group for the FIE.

As a second step, permutation testing on the inverted–upright difference wave directly probed time-point–wise differences, unconstrained by predefined temporal windows or regions of interest, providing a fuller view of Orientation effects.

Topographic maps of the difference wave revealed a negative deflection corresponding to the classic FIE—namely, a stronger N170 response to Inverted compared to Upright social stimuli, in all groups, at ∼170ms. In NA and SIB groups, this effect was lateralized to the right hemisphere, whereas in the AU group it was bilateral (Figure 4). Permutation tests supported this observation: a significant FIE was found at P8 only in the NA group, but at both P7 and P8 in the AU group. Although the SIB group exhibited a stronger FIE over the right hemisphere, it did not reach significance at either channel, which may reflect in part the smaller sample size relative to AU and NA groups. Spatio-temporal cluster analysis further revealed significant clusters over both left and right parieto-temporal sites in AU at the timing of the N170 (Figure 4C and corresponding statistical cluster plots for AU and SIB groups), whereas the inversion effects did not survive cluster-based correction for either NA or SIB groups. Additional Orientation effects in both the temporal and spatial and temporal cluster analyses can be observed in Figure 5. No effect of Orientation was observed for Objects (Supplementary Figure 1F).

**Figure 4.**
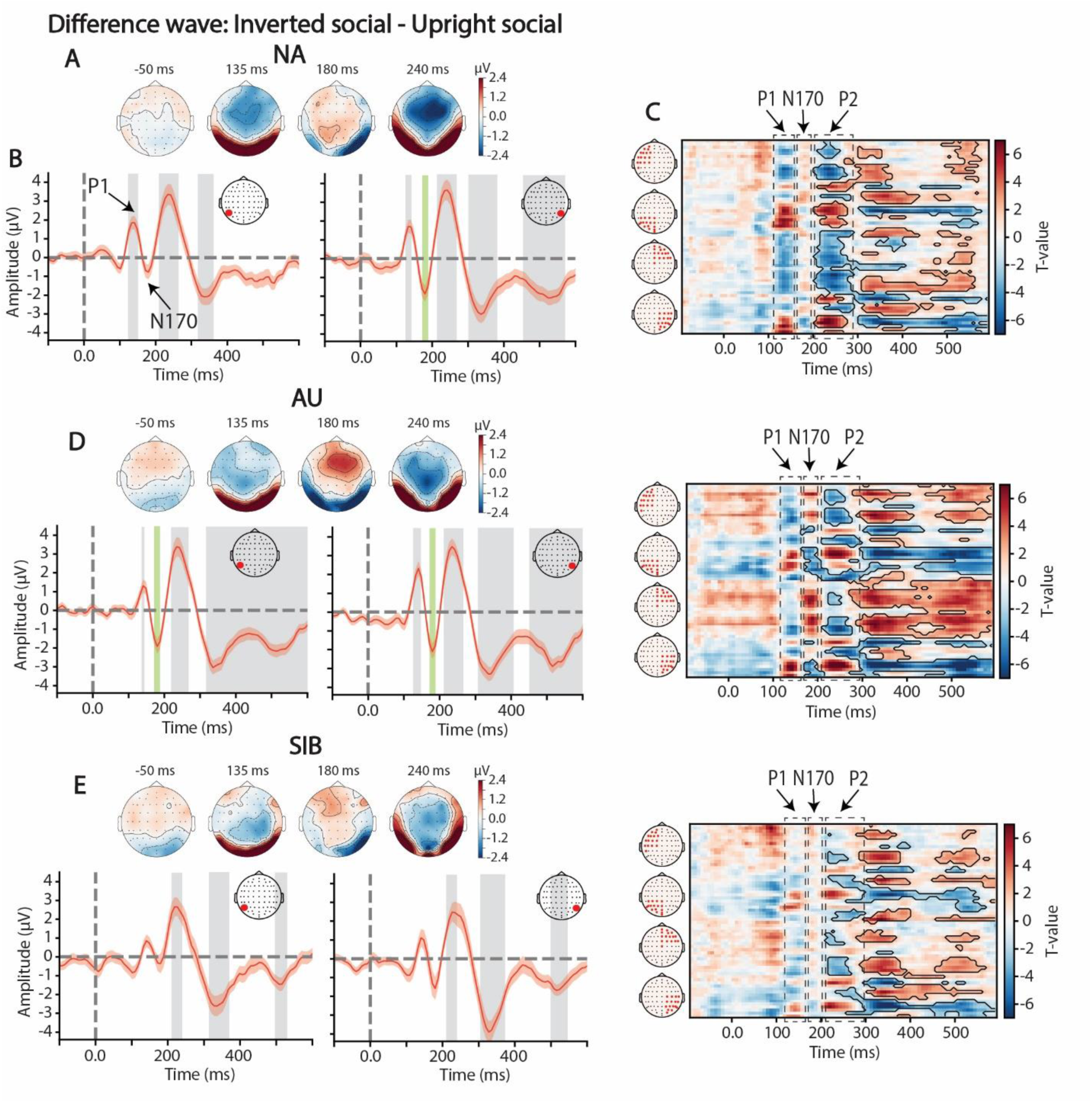
Reduced lateralization of the FIE in AU. **(A)** Topographical representation of the difference wave (Inverted social – Upright social) at the timing of the main components: P1 (135 ms), N170 (180 ms) and the P2 (240 ms) for the NA group. **(B)** Difference wave for P7 (left) and P8 (right) for the NA group. Time-points that are significantly different than 0 are highlighted in grey rectangles or a green rectangle to illustrate significant differences at the timing of the N170. **(C)** Spatio-temporal cluster plot of one-sample t-test performed for each time-point and each channel with significant spatio-temporal cluster highlighted by a black outline. **(D)** and **(E)** same structure but for AU and SIB respectively.

**Figure 5.**
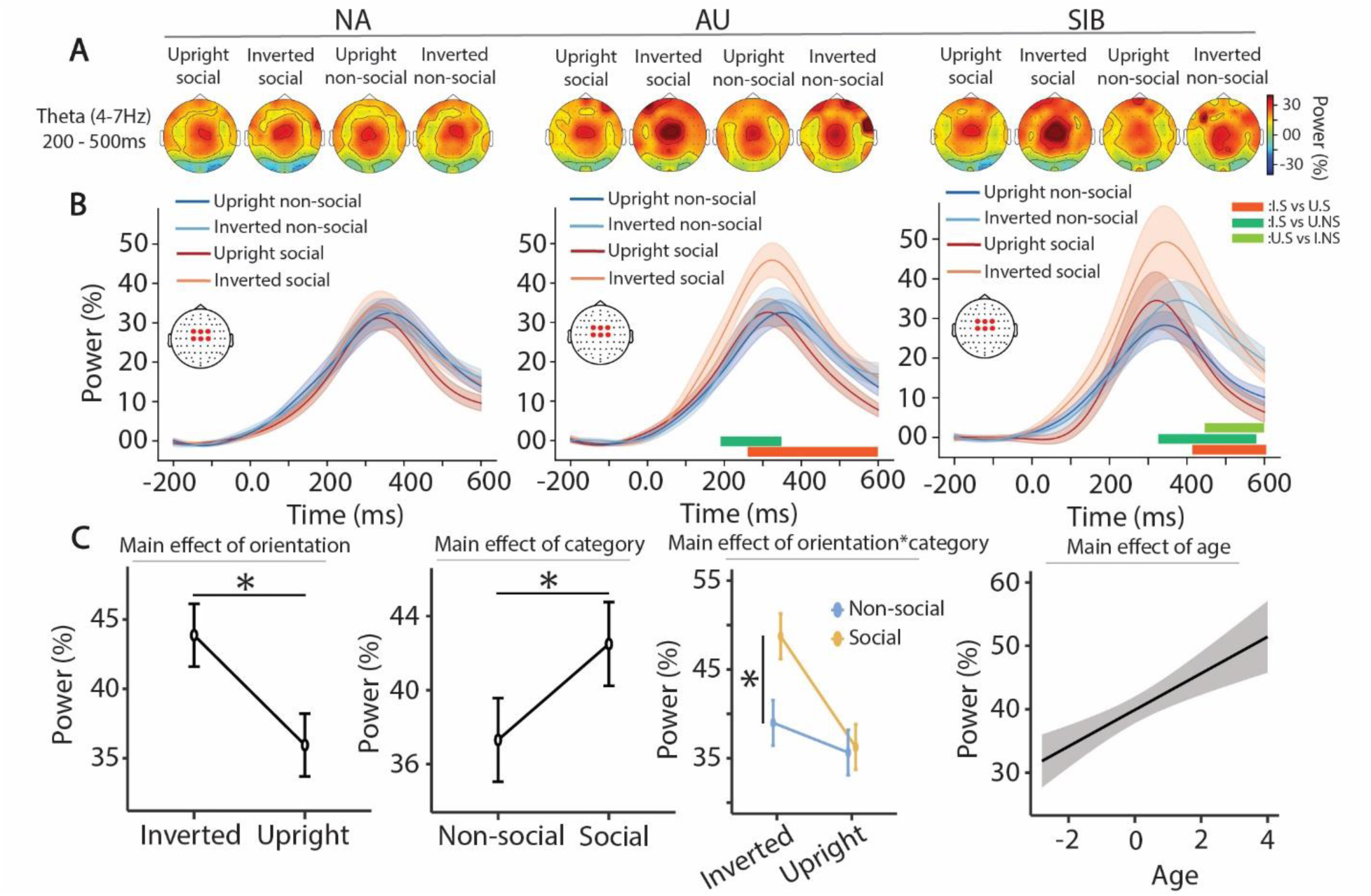
Higher theta activity for Inverted Faces in AU and SIB. **(A)** Topographical representation of the induced theta activity (4 – 7Hz) averaged between 200 to 500 ms in response to each type of stimulation and **(B)** time course of induced theta power change (in percent change from baseline) averaged over a cluster of central channels in response to Upright social (red), Inverted social (orange), Upright non-social (blue) and Inverted non-social (light blue), for NA (left), AU (middle) and SIB (right). Colored rectangles at the bottom of the plots represent paired statistical differences assessed using cluster-based permutation (orange: Upright social versus Inverted social; green: Inverted social versus Upright non-social; light green: Upright social versus Inverted non-social). **(C)** Significant main effects of the LMMs on the peak (200-500ms) of induced theta activity: Main effect of orientation; category; interaction between orientation and category and age. (*) indicates significant post-hoc (α = 0.05 and Bonferroni condition).

### Induced Theta activity in response to social and non-social stimuli

Induced theta (4-7 Hz) power was focused over central scalp for all groups (see Figure 5). While NA participants exhibited consistent levels of theta power (∼30% increase from baseline at ∼370 ms) across all conditions, statistical spatio-temporal cluster permutation analysis revealed that in AU and SIB groups, theta power was significantly greater in response to Inverted compared to Upright social stimuli (AU: 235–600 ms; SIB: 410–600 ms) and compared to Upright non-social stimuli (Inverted social vs Upright non-social: AU: 200–275 ms; SIB: 260–590 ms; Figure 5A). This difference was not evident in the NA group. To quantify this effect in a between-groups analysis, we extracted peak values from the time window (200–500 ms) at the individual level for a cluster of channels over central scalp (where the signal was maximal for all groups; Fig 5B) and submitted them to LMMs, accounting for repeated measures and including age as a covariate. LMM showed higher theta power to Inverted compared to Upright stimuli, with a main effect of Orientation (*F*(1,330) = 21.75, η²*p* = 0.06, *p* < .001; Δ = 8%) and for Social stimulation compared to Non-social with an effect of Category (*F*(1,330) = 9.34, η²*p* = 0.03, *p* = .002; Δ = 5%). A significant Category by Orientation interaction (*F*(1,330) = 7.27, η²*p* = 0.022, *p* = .007) reflected that Inverted social elicited higher theta activity than all other conditions (Upright social (Δ = 12%, *t_330_* = 5.20, *p* < .001), Inverted non-social (Δ = 10%, *t_330_* = 4.06, *p* < .001), and Upright non-social (Δ = 13%, *t_330_* = 5.45, *p* < .001)). Age was significantly associated with theta activity, with older participants showing higher theta activity (Figure 5C). The Group factor was not significant (*p* > .80).

Induced alpha desynchronization (ERD) was maximal over occipital scalp, with no significant differences across stimulus types (Condition or Orientation) or participant Group in either amplitude or topography (Supplementary Figure 2).

### Gamma-band ITPC in response to social and non-social stimuli

We next investigated inter-trial phase coherence (ITPC) in the low gamma band (25-40Hz) which has been implicated in perceptual binding and face processing (Guntekin & Basar, 2014). Prior MEG work reported reduced gamma-band ITPC in autistic individuals in response to faces, with peaks at ∼100 ms and ∼300 ms (Sun et al., 2012). Based on this, we examined group and condition differences in gamma-band ITPC across our stimuli.

Topographies revealed robust gamma ITPC increases over midline occipital electrodes (Figure 6A), with two peaks, one near 100 ms and another around 300 ms, regardless of condition or group (Figure 6B). Permutation statistics identified significant condition differences in the NA group at the first peak (100 ms), with higher gamma ITPC for both Upright and Inverted social compared to Upright and Inverted non-social. No such differences were observed in the AU group, while the SIB group showed an intermediate profile, with a significant difference only between Upright social and Upright non-social. Gamma ITPC values extracted from the first time-window (0–200 ms) at the individual level were submitted to LMMs, accounting for repeated measures and including age as a covariate. The model revealed a significant main effect of Category (*F*(1,330) = 46.62, η²p = 0.12, *p* < .001), with significantly higher gamma ITPC in response to social compared to non-social stimuli (Δ = 0.026, *t_330_* = 6.83, *p* < .001). A Group by Category interaction was also significant (*F*(2,330) = 5.39, η²*p* = 0.03, *p* = .005), with gamma ITPC higher to social than to non-social stimuli for the NA group only (Δ = 0.04, *t_330_* = 6.53, *p* < .001). These findings suggest that while early gamma-phase synchronization is present in all groups, its selectivity to social stimuli is only evident in NA (Figure 6C). Concerning theta and alpha ITPC, there is an increase in ITPC for all groups, without significant main effects (Supplementary figure 3).

**Figure 6.**
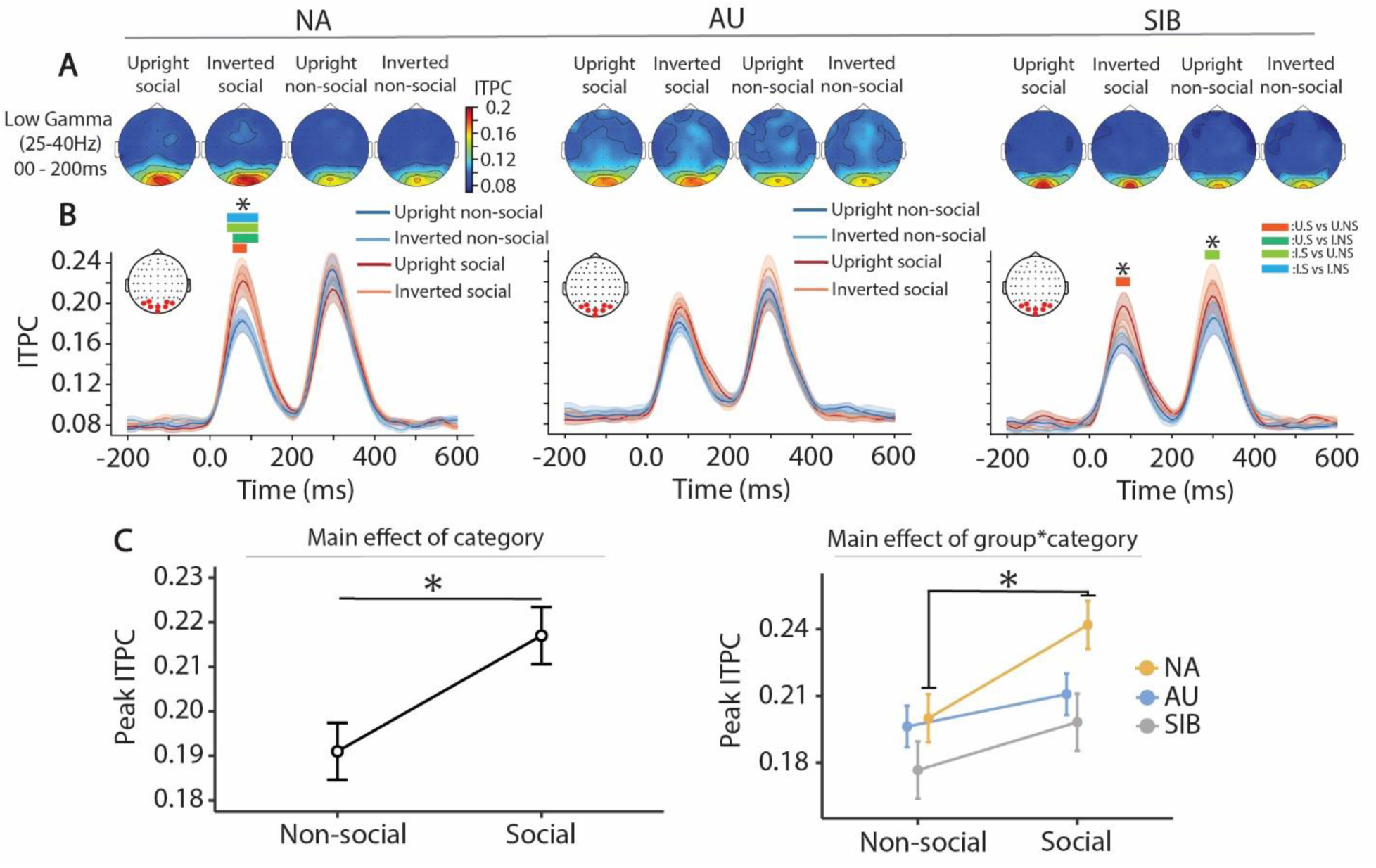
Absence of gamma ITPC selectivity to faces in AU. **(A)** Topographical representation of the low gamma (25–40Hz) ITPC averaged between 0 to 200 ms in response to each type of stimulation and **(B)** time course of the ITPC averaged over a cluster of occipital channels in response to Upright social (red), Inverted social (orange), Upright non-social (blue) and Inverted non-social (light blue), for NA (left), AU (middle) and SIB (right). Colored rectangle above the plots represents paired statistical differences assessed using cluster-based permutation (orange: Upright social versus Upright non-social; green: Upright social versus Inverted non-social; light green: Inverted social versus Upright non-social and blue: Inverted social versus Inverted non-social). **(C)** Significant main effects of the LMMs on the first peak (0-200ms) of gamma ITPC: Main effect of Category; interaction between Group and Category. (*) indicates significant post-hoc (α = 0.05 and Bonferroni condition).

## Discussion

### ERP markers of face perception in autism

The current findings demonstrate that early ERP markers of face perception (Colombatto & McCarthy, 2017; Itier et al., 2011; Olivares et al., 2015), particularly the P1 and N170, are robust across groups. Both components showed clear face selectivity, with earlier P1 latencies and larger N170 amplitudes to Social than Non-social, as well as the expected N170 enhancement for Inverted social (the face inversion effect, FIE). These results indicate that fundamental processing in sensory cortices supporting structural face encoding is typical in autistic children and their unaffected siblings, consistent with prior work reporting no differences in face processing in autism (Aydin et al., 2023; Feuerriegel et al., 2015; Shephard et al., 2020); but see (Deng et al., 2025b).

### Reduced Lateralization of the FIE in autism

Although the FIE was present in all groups, within-group comparison revealed a lack of the typical right-hemisphere dominance for the autistic group. Rather than reflecting an absence of the canonical FIE effect, this points to a deviation in the neural organization of face processing networks. Right-lateralized occipito-temporal responses are thought to support holistic face processing (Duchaine & Yovel, 2015; Rossion, 2014). The more bilateral pattern observed in AU may therefore indicate atypical hemispheric specialization or compensatory recruitment of left-hemisphere resources for face processing (Keehn et al., 2015; Li et al., 2023). Is it also notable that this finding conflicts with prior research in which the FIE was smaller for autistic individuals (Griffin et al., 2023). Although the FIE did not reach statistical significance within the SIB group, scalp topographies nonetheless suggest a qualitatively right-lateralized pattern over occipito-temporal sites (see topographical maps). This trend is consistent with a partial expression of the canonical right-hemisphere dominance for faces, albeit weaker or more variable than in NA. It is noteworthy that prior work from our group also identified highly atypical hemispheric lateralization in visual ventral stream regions in AU during an object categorization task, although in that study the stimulus materials did not include faces (Fiebelkorn et al., 2013).

Lateralization of cortical function has been documented across multiple functional domains in humans (Fair et al., 2007; Gunturkun et al., 2020; Samara & Tsangaris, 2011) including face processing (Behrmann, 2025; Behrmann et al., 2016) and is frequently diminished or atypical in neurodevelopmental and neuropsychiatric conditions (Berretz et al., 2020; Bishop, 2013; Cotter et al., 2023; de Guibert et al., 2011; Groen et al., 2008; Qi et al., 2019; Ribolsi et al., 2009; Wexler, 1980). Moreover, alterations in cortical network asymmetry have been reported in infants at elevated likelihood for autism (Rolison et al., 2022) as well as in sensory processing regions in infants later diagnosed with autism (Lewis et al., 2017). To our knowledge, our findings complement this literature by demonstrating for the first time reduced right-hemisphere dominance specifically within the face inversion context, lending further support for atypical hemispheric specialization of face-processing circuitry in autistic children.

### Differences Emerge in Oscillatory Signatures of Face Processing

While the P1 and N1 ERP components point to similar broadband sensory processing, group differences emerged in distinct oscillatory dynamics. Autistic children and their siblings showed increased frontal theta activity for Inverted social compared to Upright social stimuli, a response pattern that was absent in NA children. This suggests greater cognitive effort (Cavanagh & Frank, 2014) or compensatory control during inverted face processing conditions, mirroring findings from a previous eye-tracking study which found greater dilation to inverted faces in young autistic children as compared to their NA peers (Falck-Ytter, 2008). Gamma-band inter-trial phase coherence (∼100 ms), commonly interpreted to reflect perceptual binding, only showed face selectivity in the NA group. The level of gamma synchronization in early visual cortex represent the efficiency of the neural response and is modulated by both bottom-up stimulus properties (Friedman-Hill et al., 2000; Koelewijn et al., 2011) and top-down attentional process (Fries et al., 2002; Koelewijn et al., 2013). Intracranial and MEG studies indicate that posterior occipital regions generate face-selective gamma responses around 100 ms (Sato et al., 2014; Sun et al., 2012), and that this activity is further modulated by inversion, consistent with roles in both feature-level and configural processing (Sato et al., 2014). The absence of enhanced early gamma coherence to faces in autistic children and in first-degree relatives may thus reflect atypical binding within visual face-processing circuits (Sun et al., 2012). Consistent with prior reports of atypical gamma synchronization in autism and in relatives (Rojas et al., 2008; Sun et al., 2012; Webb et al., 2023), our results extend this literature by showing that, for upright and inverted faces, autistic children exhibit reduced gamma synchronization.

### Proposed oscillatory model for face ERPs and differences in autism

Based on the current findings of altered theta and gamma activity in the context of face processing and findings in the literature, we propose a two-stage oscillatory account of the face response. First, stimulus onset triggers a broad theta phase-reset (indexed by the robust rise in theta ITPC observed across groups and categories, see Supplementary figure 3) providing a common temporal scaffold for subsequent processing and for long range fronto-parietal communication (Fiebelkorn et al., 2018; Schnitzler & Gross, 2005; von Stein & Sarnthein, 2000). Second, theta–gamma phase–amplitude coupling (PAC) rides on this scaffold: near ∼100 ms, a gamma burst reflects local activation in occipito-temporal visual cortex (Hoogenboom et al., 2006; Muthukumaraswamy et al., 2010). In NA children, this burst becomes synchronized for social stimuli (faces), suggesting a stimulus-selective efficiency in coordinating local assemblies. For autistic children, this face-selective synchrony fails to materialize (with SIBs intermediate), consistent with diminished or more variable coupling of local populations to the theta scaffold for socially salient inputs. The observation of a second gamma peak ∼200 ms after the first further supports a PAC account, aligning with ∼5 Hz theta gating (Lisman & Jensen, 2013).

In sum, while broadband ERPs (P1/N170) appear similar across groups, an oscillatory decomposition reveals reduced early network synchronization to faces in autistic children. Interestingly, another study found atypical coupling of gamma and theta activity during speech processing in autism (Jochaut et al., 2015); autistic individuals exhibited a lack of down-regulation of gamma activity by theta during speech stimuli presentation. This atypical dependency disrupts the delicate balance crucial for accurate auditory processing, potentially leading to difficulties in speech decoding and communication. Such findings suggest that atypical coordination of cortical oscillations may underlie the characteristic social communication difficulties observed in autism, whether in the auditory or visual domain.

## Limitations

This study should be considered in light of several limitations. First, the sample was restricted to children aged 8–13 years, which constrains the generalizability of findings to other developmental stages, and group sizes for SIB and NA participants were relatively modest. Second, the task design emphasized positive-emotion faces and a low-level target detection requirement, limiting conclusions about higher-order aspects of social cognition including emotion processing.

Third, although our results motivate a theta–gamma coupling account, the present scalp EEG data are not well suited to directly test PAC. High-frequency activity is substantially attenuated and spatially blurred by the scalp and skull, reducing single-trial signal-to-noise for gamma bursts. In addition, the very high theta ITPC we observe (while central to our model) also reduces across-trial phase variability, which undermines classical PAC estimators that rely on sufficient dispersion of theta phase and gamma amplitude across trials. Finally, robust tests of burst timing relative to theta phase (rather than averaged power) require reliable single-trial gamma burst detection, which scalp EEG at our sampling rate and SNR cannot provide. Consequently, we treat the PAC account as a plausible, testable framework to be examined in future work, not a confirmed mechanism.

Finally, our sample consisted of high-functioning autistic children, representing only a subset of the autism spectrum. These findings may not generalize to individuals with lower cognitive or language abilities, and future studies should explore whether similar mechanisms are present across a broader range of functioning levels.

## Implications

Together, these findings suggest a two-level framework for understanding early face processing in autism: (1) broad-band sensory-driven mechanisms for face perception (P1/N170) remain robust in autistic children (Aydin et al., 2023; Feuerriegel et al., 2015; Naumann et al., 2018; Shephard et al., 2020), supporting similar bottom-up sensory driven time locked encoding, while (2) specific oscillatory processes involving network-level coordination show atypical modulation. Notably, we observed these oscillatory differences in a very simple paradigm that required no identity or emotion judgments. We therefore expect these subtle divergences to become more consequential in more complex, naturalistic settings, where functions subserved by these oscillations (e.g., binding, attention, control) are more heavily taxed.

These findings also add nuance to ongoing debates about candidate biomarkers. For instance, N170 latency has been proposed as a potential biomarker of autism (Kala et al., 2021), yet our results revealed no group effects on latency under simple conditions. This suggests that latency measures may lack sensitivity in basic paradigms, whereas oscillatory indices of network-level dynamics may provide richer biomarkers of atypical processing trajectories. For instance, future work should examine cross-frequency interactions between theta and prioritize multimodal approaches, combining ERP and oscillatory measures with behavioral and gaze-based indices, to better characterize the mechanistic pathways underlying social perceptual differences in autism.

## Conclusions

In sum, our findings indicate that while stimulus driven encoding of faces is broadly typical in autistic children, atypicalities emerge in the coordination and specialization (lateralization) of neural networks. This dissociation suggests that difficulties in holistic and socially meaningful face perception for autistic children arise not from failure to detect or encode faces, but from inefficiencies in integrating these signals within broader perceptual and attentional systems. By highlighting where processing occurs similarly and where it diverges, this work refines our understanding of the neural architecture of face perception in autism and underscores the value of moving beyond single-component ERP markers toward oscillatory and network-level measures. Such approaches may better capture the dynamic, context-dependent processes most relevant for identifying mechanistic biomarkers and informing interventions that target social perception.

## Declarations

Ethics approval and consent to participate: This study was approved by the Institutional Review Board of the Albert Einstein College of Medicine (IRB # 2021-13433). All participants assented to the procedures and parents/guardians provided informed consent.

Availability of data and materials: The dataset supporting the conclusions of this article is available in the ‘SFARI_EEG_multi-paradigm dataset’ (‘FAST’ paradigm) repository (BIDS format), doi:10.18112/openneuro.ds006780.v1.0.0. The scripts used for preprocessing and analyses of the data are available on GitHub: https://github.com/tvanneau/SFARI-FAST. Competing interest: The authors declare no competing interests.

Author contributions: S.M conceived the study. T.V , C.B, and M.D preprocessed the data. T.V and C.B analyzed the data under the supervision of S.M . T.V and C.B wrote the first draft of the manuscript. J.J.F, S.M, and M.D edited the manuscript.

## Acknowledgments

The authors thank Daniella Coen, Albulena Sejdiu, Dennis Cregin and Tringa Lecaj for their help with experiments. We are grateful to the families and all the individuals that participated in the study.

Funding: This work was supported by a grant from the Simons Foundation Autism Research Initiative (SFARI Award # 874845, SM); Support for recruitment and phenotyping of participants was provided by the Human Clinical Phenotyping Core of the NICHD-funded Rose F. Kennedy Intellectual and Developmental Disabilities Research Center (P50 HD105352 – to SM). Work at Rochester is supported through the Golisano Intellectual and Developmental Disabilities Research Institute (UR-IDDRC), which is supported by a center grant from the Eunice Kennedy Shriver National Institute of Child Health and Human Development (P50 HD103536 – to JJF). The content is solely the responsibility of the authors and does not necessarily represent the official views of the National Institutes of Health.

## List of abbreviations

ADI-R: (Autism Diagnostic Interview–Revised)
ADOS-2: (Autism Diagnostic Observation Schedule Second Edition)
ANOVA: (Analysis of Variance)
AU: (Autistic group)
BAP: (Broad Autism Phenotype)
CARS-2: (Childhood Autism Rating Scale Second Edition)
CMS/DRL: (Common-Mode-Sense/Drive-Right-Leg the BioSemi reference system)
EEG: (Electroencephalography)
ERD: (Event-Related Desynchronization)
ERP: (Event-Related Potential)
FIE: (Face Inversion Effect)
FIR: (Finite Impulse Response)
FS-IQ: (Full-Scale Intelligence Quotient)
ICA: (Independent Component Analysis)
IDD: (Intellectual and Developmental Disability)
IRB: (Institutional Review Board)
ITPC: (Inter-Trial Phase Coherence)
LMM: (Linear Mixed Model)
MEG: (Magnetoencephalography)
MNE: (Magnetoencephalography and EEG Python analysis toolbox)
NA: (Non-Autistic group)
PAC: (Phase–Amplitude Coupling)
ROI: (Region of Interest)
RT: (Reaction Time)
SIB: (Sibling group)
SNR: (Signal-to-Noise Ratio)
VEP: (Visual Evoked Potential)

---

**Supplementary figure 1.**
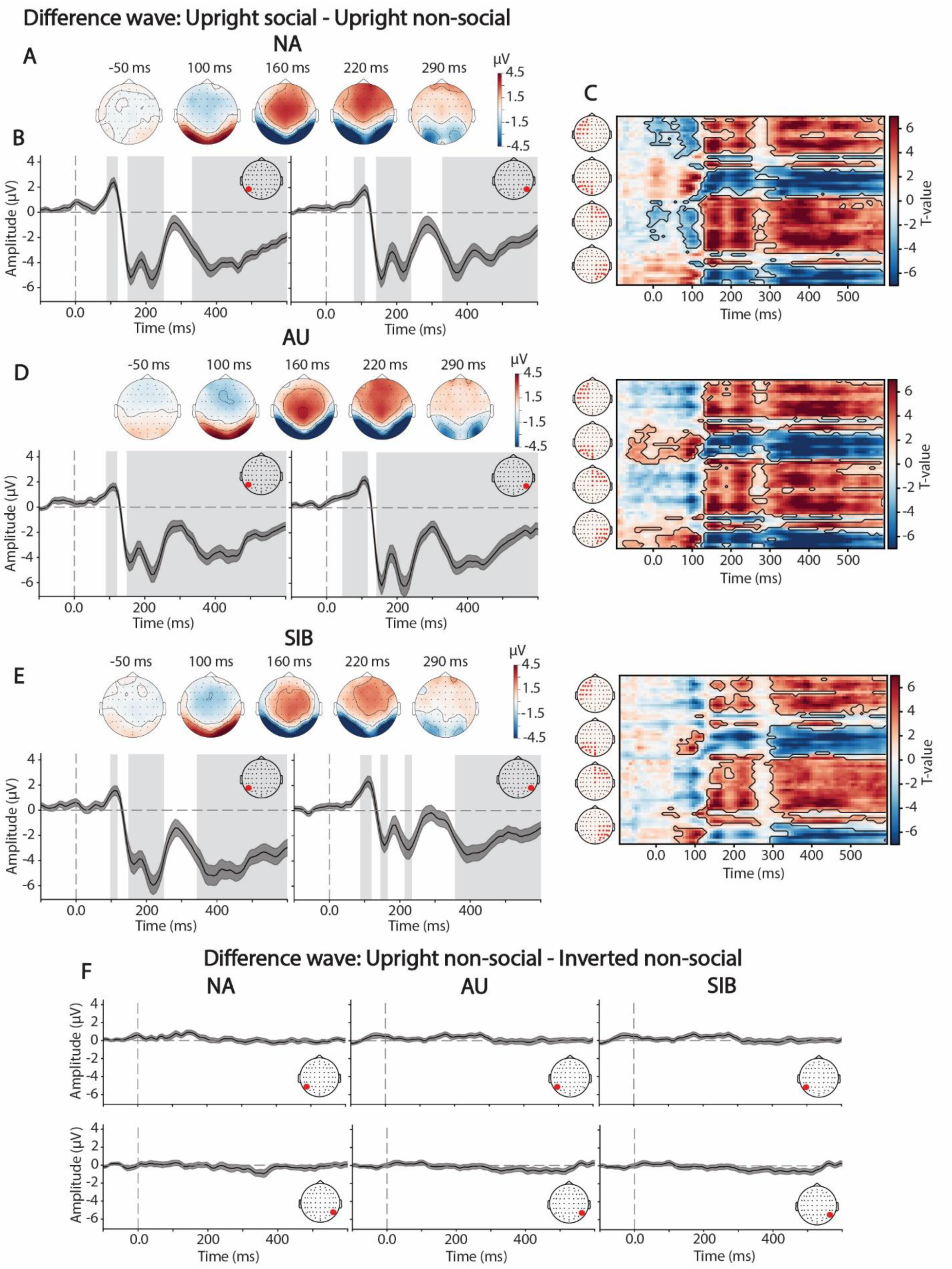
**(A)** Topographical representation of the difference wave (Upright social – Upright non-social) at the timing of the main components: P1 (135 ms), N170 (180 ms) and the P2 (240 ms) for the NA group. **(B)** Difference wave for P7 (left) and P8 (right) for the NA group. Time-points that are significantly different than 0 are highlighted in grey rectangles or a green rectangle to illustrate significant differences at the timing of the N170. **(C)** Spatio-temporal cluster plot of one-sample t-test performed for each time-point and each channel with significant spatio-temporal cluster highlighted by a black outline. **(D)** and **(E)** same structure but for AU and SIB respectively. (F) Difference wave (Upright non-social – Inverted non-social) for NA (left), AU (middle) and SIB (right) groups, for P7 (top row) and P8 (bottom row).

**Supplementary figure 2.**
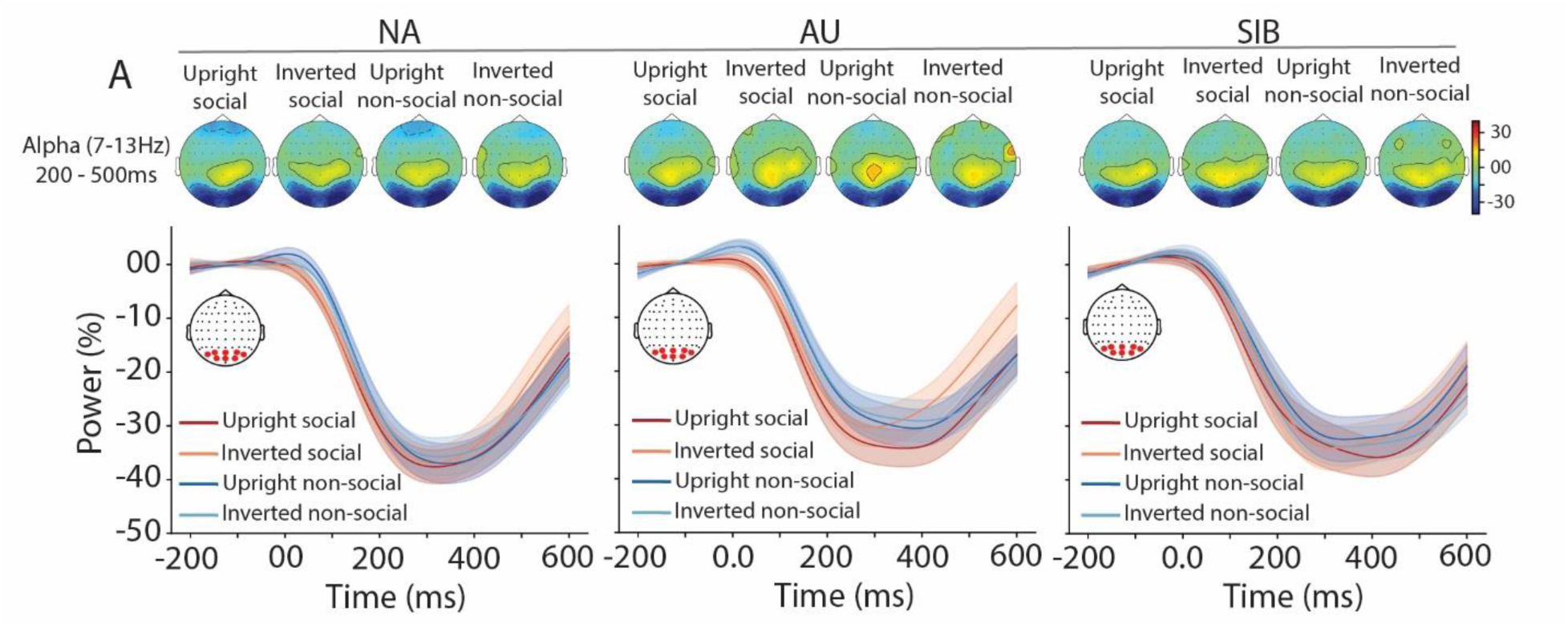
Similar alpha event-related desynchronization between groups and conditions. **(A)** Topographical representation of the induced alpha activity (7 – 13Hz) averaged between 200 to 500 ms in response to each type of stimulation and **(B)** time course of induced alpha power change (in percent change from baseline) averaged over a cluster of occipital channels in response to Upright social (red), Inverted social (orange), Upright non-social (blue) and Inverted non-social (light blue), for NA (left), AU (middle) and SIB (right)., for NA (left), AU (middle) and SIB (right).

**Supplementary figure 3.**
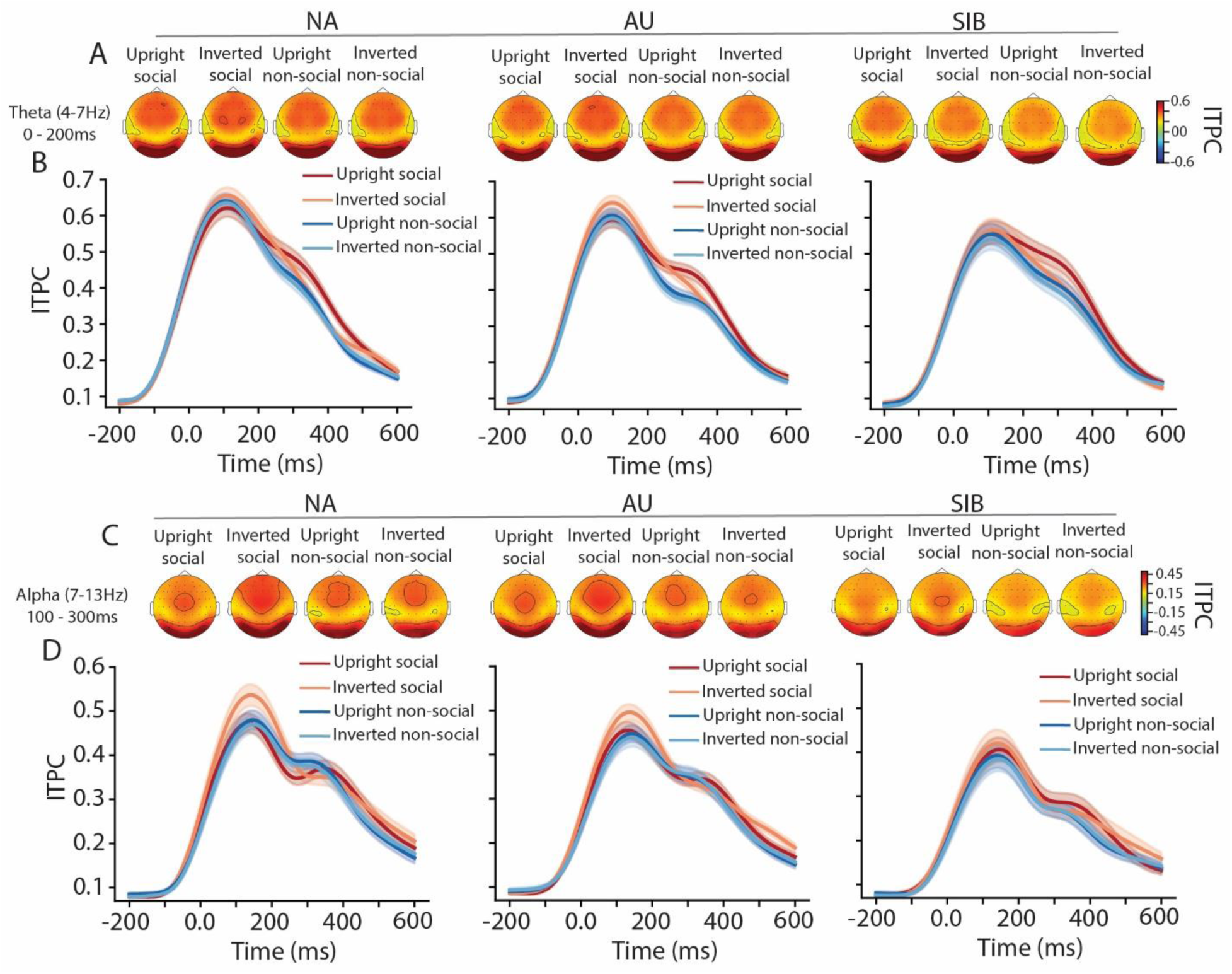
Inter-trial phase coherence (ITPC) for theta and alpha band. **(A)** Topographical representation of the ITPC for the theta band (4 – 7Hz) averaged between 0 to 200 ms in response to each type of stimulation and **(B)** time course of the ITPC averaged over a cluster of occipital channels in response to Upright social (red), Inverted social (orange), Upright non-social (blue) and Inverted non-social (light blue), for NA (left), AU (middle) and SIB (right). (**C**) and (**D**) same as (**A**) and (**B**) but for alpha (7 – 13Hz) activity.

